# Accumulation of sensory evidence is impaired in Parkinson’s disease with visual hallucinations

**DOI:** 10.1101/111278

**Authors:** Claire O’Callaghan, Julie M. Hall, Alessandro Tomassini, Alana J. Muller, Ishan C. Walpola, Ahmed A. Moustafa, James M. Shine, Simon J. G. Lewis

**Author notes:** Corresponding author: Claire O’Callaghan, Herchel Smith Building for Brain & Mind Sciences, Cambridge Biomedical Campus, Cambridge CB2 0SZ, UK.

## Abstract

Models of hallucinations across disorders emphasise an imbalance between sensory input and top-down influences over perception. However, the psychological and mechanistic correlates of this imbalance remain underspecified. Visual hallucinations in Parkinson’s disease (PD) are associated with impairments in lower level visual processes and attention, accompanied by over activity and connectivity in higher-order association brain networks. PD therefore provides an attractive framework to explore the relative contributions of bottom-up versus top-down disturbances in hallucinations. Here, we characterised sensory processing in PD patients with and without visual hallucinations, and in healthy controls, by fitting a hierarchical drift diffusion model (hDDM) to an attentional task. The hDDM uses Bayesian estimates to decompose reaction time and response output into parameters reflecting drift rates of evidence accumulation, decision thresholds and non-decision time. We observed slower drift rates in PD patients with hallucinations, which were insensitive to changes in task demand. In contrast, wider decision boundaries and shorter non-decision times relative to controls were found in PD regardless of hallucinator status. Inefficient and less flexible sensory evidence accumulation emerge as unique features of PD hallucinators. We integrate these results with current models of hallucinations, suggesting that slow and inefficient sensory input in PD is less informative, and may therefore be down-weighted leading to an over reliance on top-down influences. Our findings provide a novel computational framework to better specify the impairments in dynamic sensory processing that are a risk factor for visual hallucinations.

---

Visual hallucinations are common in Parkinson’s disease (PD), occurring in over 30% of newly diagnosed and early-stage patients, and increasing to upwards of 70% by the late stages of the disease (Holroyd *et al.*, 2001, Hely *et al.*, 2008, Pagonabarraga *et al.*, 2016). But despite their prevalence, visual hallucinations remain poorly understood and treatment options are limited (ffytche *et al.*, 2017). Continued characterisation of the psychological and mechanistic correlates of visual hallucinations in PD will be crucial to inform therapeutic advances.

Proposed explanatory models for visual hallucinations in PD emphasise a state of reduced sensory input, where the ongoing perceptual process is vulnerable to influence from internally generated imagery (Hobson *et al.*, 2000, Collerton *et al.*, 2005, Diederich *et al.*, 2005, Shine *et al.*, 2014). This is in keeping with a transdiagnostic framework, where hallucinations arise when the balance between sensory input and top-town influence over perception is disrupted, such that sensory information is reduced or not properly integrated and there is a predominance of top-down influence (Friston, 2005, Fletcher and Frith, 2009, Adams *et al.*, 2013, Powers *et al.*, 2016, O’Callaghan *et al.*, 2017).

In PD, sensory input is affected by dopaminergic retinal changes and impairments in lower level visual processes and attention - all of which can be more pronounced in patients with hallucinations (Weil *et al.*, 2016). However, hallucinations can also occur in patients where ophthalmological measures and performances on lower level perceptual tasks are equivalent to non-hallucinating patients (Gallagher *et al.*, 2011). A possibility is that lower level sensory impairment and reduced attention confer risk factors for visual hallucinations in PD, but failures in the dynamic integration of visual input and attention trigger their occurrence (Collerton *et al.*, 2005, Diederich *et al.*, 2005). Here, we aimed to investigate the dynamic processes underlying visual perception in PD hallucinators by applying a drift diffusion model (DDM) to an attentional task.

DDMs, like other models of perceptual decision-making, are based on the premise that reaction time and response output can be decomposed into parameters reflecting the latent cognitive processes driving task performance (Mulder *et al.*, 2014). The DDM is particularly relevant to assessing sensory information processing in PD hallucinators, as it quantifies information extracted from a stimulus (drift rate), the evidence needed to make a decision (boundary separation), and components related to stimulus encoding and response output (non-decision time) (Voss *et al.*, 2004, Ratcliff and McKoon, 2008). In this study, we applied a Bayesian hierarchical version of the DDM, which is robust in the context of low trial numbers (Wiecki *et al.*, 2013, Cavanagh *et al.*, 2014). This makes the task suitable for clinical contexts where task duration is necessarily limited, and it has been successfully fitted to data from PD patients in previous studies (Cavanagh *et al.*, 2011, Herz *et al.*, 2016b, Zhang *et al.*, 2016). We assessed participants on the attentional networks task (Fan *et al.*, 2002), which allowed us to measure perceptual decision making under conditions with different levels of difficulty as determined by perceptual conflict in the stimuli (neutral and congruent conditions vs. incongruent condition). We predicted that parameters reflecting the integration of sensory evidence in the decision making process, i.e., the drift rate or boundary separation, would be impaired in hallucinators relative to non-hallucinators, but that their non-decision components would be similar. We also predicted that controls and non-hallucinators would modulate their drift rate and boundary separation in response to the different levels of perpetual conflict, but that hallucinators would not show the same level of flexibility in response to task demands.

## Methods and Materials

### Case selection

A total of 50 patients were recruited from the Parkinson’s disease research clinic at the Brain and Mind Centre, University of Sydney, Australia. Patients were identified as hallucinators if they self-reported visual hallucinatory phenomena and scored ≥1 on question two of the MDS-UPDRS (i.e., over the past week have you seen, heard, smelled or felt things that were not really there? If yes, examiner asks the patient or caregiver to elaborate and probes for information) (Goetz *et al.*, 2008). This resulted in a group of 24 hallucinators (VH) and 26 non-hallucinators (nonVH). Four patients from the VH group and 1 from the nonVH group were excluded from analysis due to excessive missed responses on the experimental task, leaving a final cohort of 20 hallucinators and 25 non-hallucinators. A proportion of these patients were included in a previous study involving a behavioural investigation of the attentional networks task (Hall *et al.*, 2016). Twelve aged-matched controls were recruited from a volunteer panel.

All patients satisfied the United Kingdom Parkinson’s Disease Society Brain Bank criteria and were not demented (Martinez-Martin *et al.*, 2011). Patients were assessed on the Hoehn and Yahr Scale and the motor section of the Unified Parkinson’s Disease Rating Scale (UPDRS-III). The Mini-mental state examination (MMSE) and Montreal Cognitive Assessment (MoCA) were administered as measures of general cognition. Clinical assessments and the experimental task were performed with patients in the ON state having taken their regular dopaminergic medication, and dopaminergic dose equivalence scores were calculated. No patients in the cohort were taking antipsychotic medication or cholinesterase inhibitors. Control participants were screened for a history of neurological or psychiatric disorders. Patients and controls were matched for age. The study was approved by the local ethics committees and all participants provided informed consent in accordance with the Declaration of Helsinki.

### Attentional network task (ANT)

We administered a shortened version of the ANT (Fan *et al.*, 2002), which requires participants to determine if a central arrow points left or right. Central arrows are flanked by flat lines (neutral condition), arrows facing the same direction (congruent condition), or arrows facing a mixture of directions (incongruent condition) (See figure 1, Panel a). The perceptual conflict in the incongruent condition is designed to place a greater demand on attentional processes, relative to the congruent and neutral conditions. The ANT also contains spatial and warning cues within the three conditions to evaluate alerting and orienting, but these were not analysed in the current study. A total of 96 trials were administered (32 from each condition); target stimuli were displayed for a maximum of 1700 msec and responses for left and right were made using the *z* and *m* keys on a standard keyboard.

**Figure 1.**
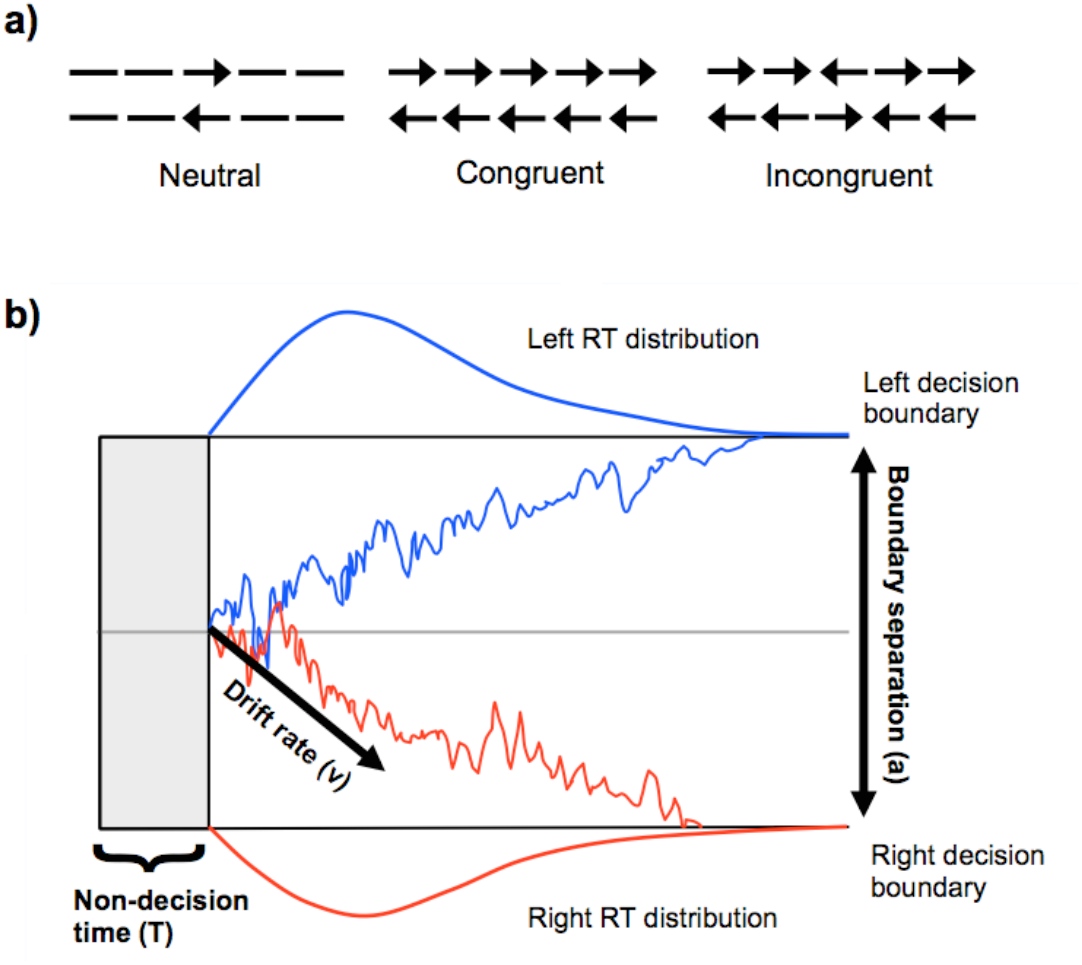
Attentional networks task conditions and drift diffusion model. ***Legend –***Panel a) The three conditions in the attentional networks task. Panel b) Example of drift diffusion trajectories. Evidence is noisily accumulated toward a left or right response (blue and red panels), which are separated by the boundary threshold (*a*). The average evidence accumulation is denoted by drift rate (*v*). The evidence accumulation begins after a period of non-decision time (*T*). Density plots show the distribution of observable reaction times (RT). Adapted from (Wiecki *et al.*, 2013, Zhang and Rowe, 2014).

### Hierarchical drift-diffusion model of the ANT

Drift-diffusion models (DDMs) are widely applied to rapid, two-choice decision making tasks such as the ANT (Ratcliff and McKoon, 2008, Wagenmakers, 2009, Ratcliff *et al.*, 2016). DDMs are typically described by four main parameters: drift rate *v*, decision boundary *a,* decision bias *z* and non-decision time *T*. The decision process is modelled as the gradual accumulation of information, reflected by the drift rate, which continues until a decision boundary is reached. Fast and accurate decisions are often generated by high drift rates, whereas lower drift rates lead to slow and error-prone decisions (Krypotos *et al.*, 2015). Separation of upper and lower decision boundaries reflects response caution, where high values are associated with accurate, long reaction times and low values with shorter and more error prone reaction times (Krypotos *et al.*, 2015). The decision bias parameter captures *a priori* bias toward one of the two responses. Non-decision time incorporates components that are not part of the evidence accumulation process, including stimulus encoding, extracting stimulus dimensions and executing a response (Ratcliff *et al.*, 2016). See Figure 1, Panel b, for an example of the drift diffusion process. In the current study we applied a hierarchical DDM (hDDM), using the hDDM toolbox (http://ski.clps.brown.edu/hddm_docs/ (Wiecki *et al.*, 2013) to fit the response and reaction time data from the ANT. The hDDM uses Bayesian estimation to generate posterior distributions of DDM parameters at group and subject levels. This approach optimises the trade-off between random and fixed effect models, accounting for both within-subject variability and group level similarities, as individual parameters are constrained by a group level distribution.

We tested three models that all assumed an unbiased starting point (*z.*, given that left/right responses were counterbalanced, and assumed that non-decision time (*T*) would not be expected to vary as a function of condition, as the stimulus encoding and motor responses required across conditions were comparable. Model specifications were as follows: in the first model, only drift rate (*v*) was permitted to vary by condition, and decision boundary (*a*) was held constant; in a second model, *a* could vary across conditions, but *v* was held constant; in a third model, both *v* and *a* were free to vary across conditions. To optimise convergence only group level parameters were estimated (Wiecki *et al.*, 2013). For all models, Markov Chain Monte Carlo simulations were used to generate 120,000 samples from the joint posterior parameter distribution. The first 20,000 samples were discarded as burn-in and we used a thinning factor of 10, with outliers specified at 5%. Convergence was assessed by visually inspecting the Markov chains and computing the R-hat Gelman-Rubin statistic where successful coverage is indicated by values <1.1 (Krypotos *et al.*, 2015). The best model was determined by comparing the deviance information criterion (DIC) of each model, which evaluates a model’s goodness-of-fit while accounting for model complexity (i.e., number of free parameters), with lower DIC values indicating better model fit (Spiegelhalter *et al.*, 2002). To further evaluate the best fitting model, we ran posterior predictive checks by averaging 500 simulations generated from the model’s posterior to confirm it could reliably reproduce patterns in the observed data (Wiecki *et al.*, 2013). The fitted reaction time, choice data and source code for the hDDM can be found at https://github.com/claireocallaghan/hDDM_ANT_PD.

### Statistical analysis

Independent samples t-tests and ANOVAs with Tukey *post hoc* tests were used to compare demographics and behavioural results from the ANT. Parameters from the hDDM were analysed using Bayesian hypothesis testing to determine the extent of overlap between the percentage of samples drawn from two posterior density distributions. Posterior probabilities are considered significantly different if less than 5% of the distributions overlap (Wiecki *et al.*, 2013, Cavanagh *et al.*, 2014, Herz *et al.*, 2016b). Here, we applied Bonferroni correction for multiple comparisons (5% /13) where significance is assigned when less than 0.38% of the posterior distributions overlap. The proportion of overlap in the posterior probabilities is denoted by *P* to distinguish it from the classical frequentist *p* values.

## Results

### Participant characteristics

Demographic and clinical characteristics are shown in Table 1. The groups were matched for age (F(2,54) = 1.31, *p* = 0.28). Performance on the MMSE was similar across groups (F(2,54) = 1.07, *p* = 0.35), but the MoCA revealed significant group differences, with the nonVH group performing below control levels [(F(2,54) = 5.91, *p* < 0.01); nonVH vs. controls: p<0.01; VH vs controls and nonVH: *p* values = 0.17 and 0.19)]. The patient groups did not differ in disease duration [*t*(−1.42), *p* = 0.16], Hoehn and Yahr stage [*t*(−1.33), *p* = 0.19], UPDRS III [*t*(−1.80), *p* = 0.08], or dopamine dose equivalence [*t*(-1.63), *p* = 0.11].

**Table 1.** Demographics and clinical characteristics of Parkinson’s disease and healthy control participants.

|  | Control | PD nonVH | PD VH |
| --- | --- | --- | --- |
| **N** | 12 | 25 | 20 |
| **Sex (M:F)** | 4:8 | 19:6 | 15:5 |
| **Age** | 64.75 (5.97) | 67.08 (8.22) | 68.89 (6.05) |
| **MMSE** | 29.08 (.79) | 28.39 (1.47) | 28.79 (1.72) |
| **MoCA** | 28.50 (1.45) | 26.56 (2.11) | 27.41 (1.50) |
| **Disease duration (yrs)** | - | 5.52 (2.86) | 7.11 (4.99) |
| **Hoehn & Yahr stage** | - | 2.16 (.47) | 2.37 (.60) |
| **UPDRS III** | - | 27.67 (11.54) | 34.28 (13.97) |

Values are mean (standard deviation).

### ANT Behavioural results

For the ANT, participants who made no response on more than one third of trials were excluded from the study. This resulted in the exclusion of one nonVH and 4 VH patients. Trials where no response was made were omitted from the behavioural and modelling analyses, rather then using the upper limit reaction time, which would bias the model. After removal of no response trials, accuracy was at 100% across the three groups. Reaction times are plotted in Figure 2. Global reaction times, regardless of condition, were fastest for controls, followed by nonVH, then VH, with significant differences evidenced by a main effect for group in the ANOVA [F(2,162) = 8.15, *p* < 0.001; VH vs controls and nonVH: *p* values < 0.01; nonVH vs controls: *p* = 0.59]. A main effect of condition revealed that reaction times were significantly slower for the incongruent condition compared to both the congruent and neutral conditions, whereas the congruent and neutral conductions were equivalent [F(2,162) = 17.02, *p* < 0.000001; incongruent vs congruent and neutral: *p* values < 0.00001; congruent vs neutral: *p* = .98]. There was no significant interaction between group and condition, suggesting that the relatively slowed reaction times for the incongruent condition were consistent across groups [F(4,162) = .04, *p* = .99].

**Figure 2.**
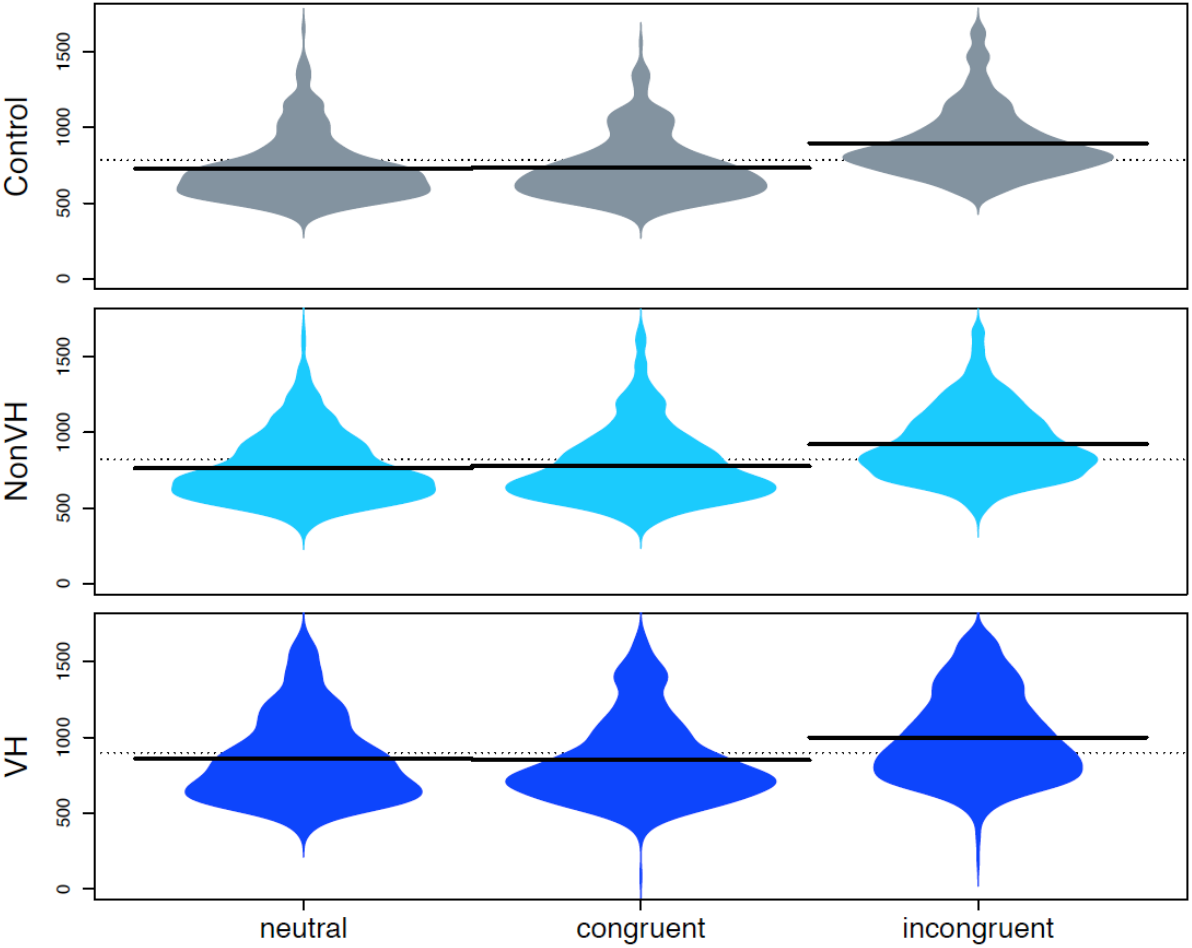
Attentional networks task reaction times. ***Legend –***Reaction time distributions for the attentional networks task across the three levels of task difficulty. Bolded black lines designate mean reaction times.

### Hierarchical drift-diffusion model fit

All three models showed good convergence, based on R-hat values under 1.1 and visually inspected chains (see supplementary material for R-hat values). The best fitting model was model three, which allowed *v* and *a* to vary across conditions (DIC model 3: −636.496; compared to DIC model 1: −436.160 and DIC model 2: −626.593). Posterior predictive checks showed good agreement between the simulated and observed data as shown in supplementary figure 1 plotting the observed data against the model prediction. Comparisons showed that the difference between the summary statistics of the simulated and the observed data fell within the 95% credible interval.

### Analysis of model parameters

#### Comparisons between groups

Figure 3 shows the posterior probability density plots for the drift rate (top panel) and decision boundaries (bottom panel) for the three groups across each condition. The VH group had uniformly lower drift rates compared to nonVH, these differed significantly in the neutral and incongruent conditions (P = 0.05% and *P* = 0.04%) but not in the congruent condition *(P* = 3.41%). Drift rates of VH patients were also significantly lower than controls for the incongruent condition (P = 0.04%), although not for the neutral and congruent conditions (P = 3.43% and *P* = 12.72%). Posterior probabilities did not differ significantly between nonVH and control groups for any condition (neutral: *P* = 84.21%; congruent: *P* = 66.45%; incongruent: *P* = 41.19%).

**Figure 3.**
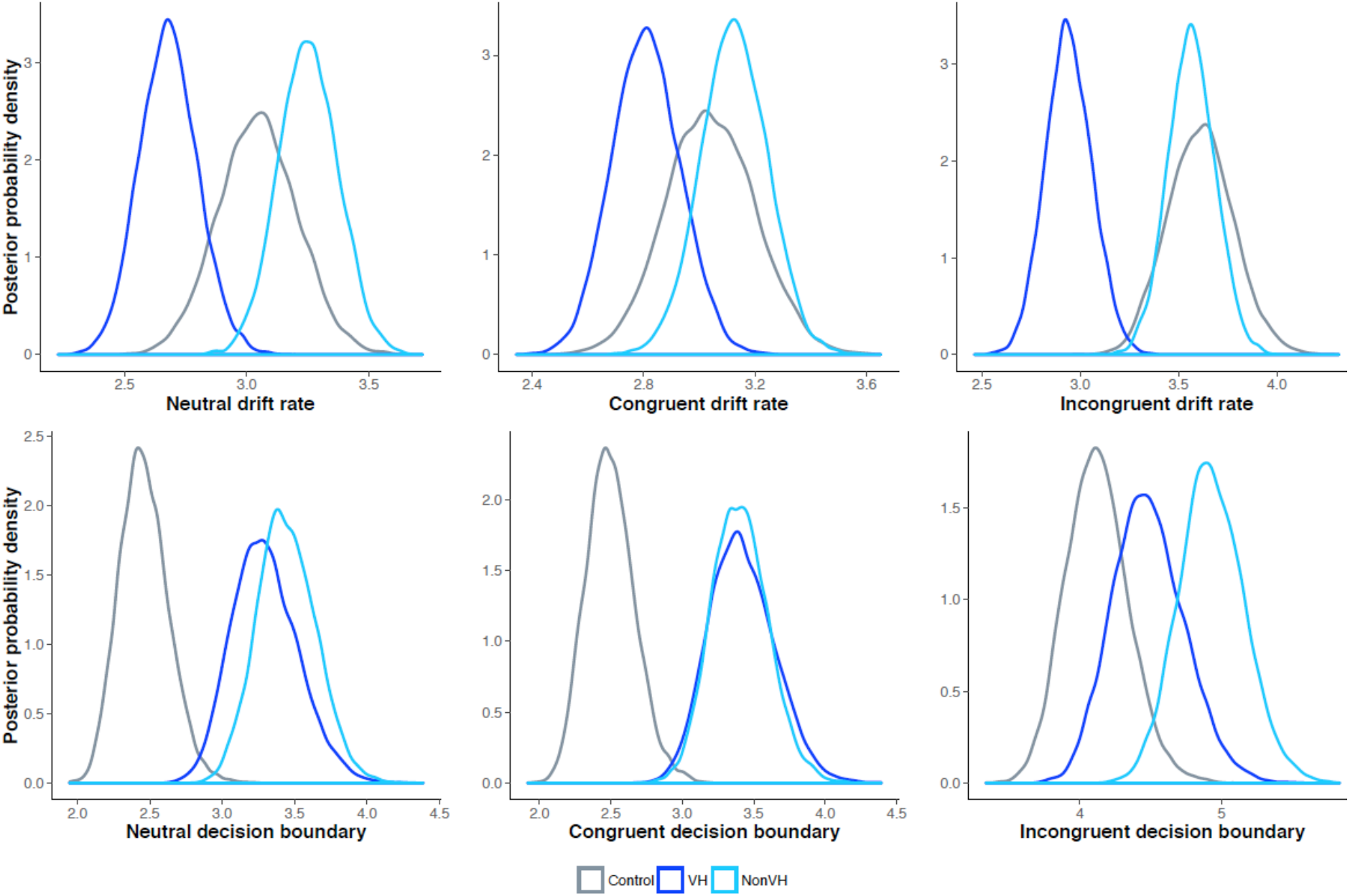
Between group comparisons of drift rates and decision boundaries. ***Legend –*** Posterior probability density plots for drift rates (top panel) and decision boundaries (bottom panel).

For the decision boundary, there were no significant differences between the VH and nonVH groups for any of the conditions (neutral: *P* = 31.19%; congruent: *P* = 50.96%; incongruent: *P* = 9.45%). Both the VH and nonVH group had significantly larger decision boundaries than controls in the neutral and congruent conditions (VH: *P* = 0.12% and *P* = 0.03%; nonVH: *P* = 0.03% and *P* = 0.03%), but not the incongruent condition (VH: *P* = 14.19% nonVH: *P* = 0.68%).

As shown in figure 4, VH and nonVH groups had similar non-decision times (*P* = 34.04%), which were significantly shorter than controls (VH: *P* = 0.05% nonVH: *P* = 0.00%).

**Figure 4.**
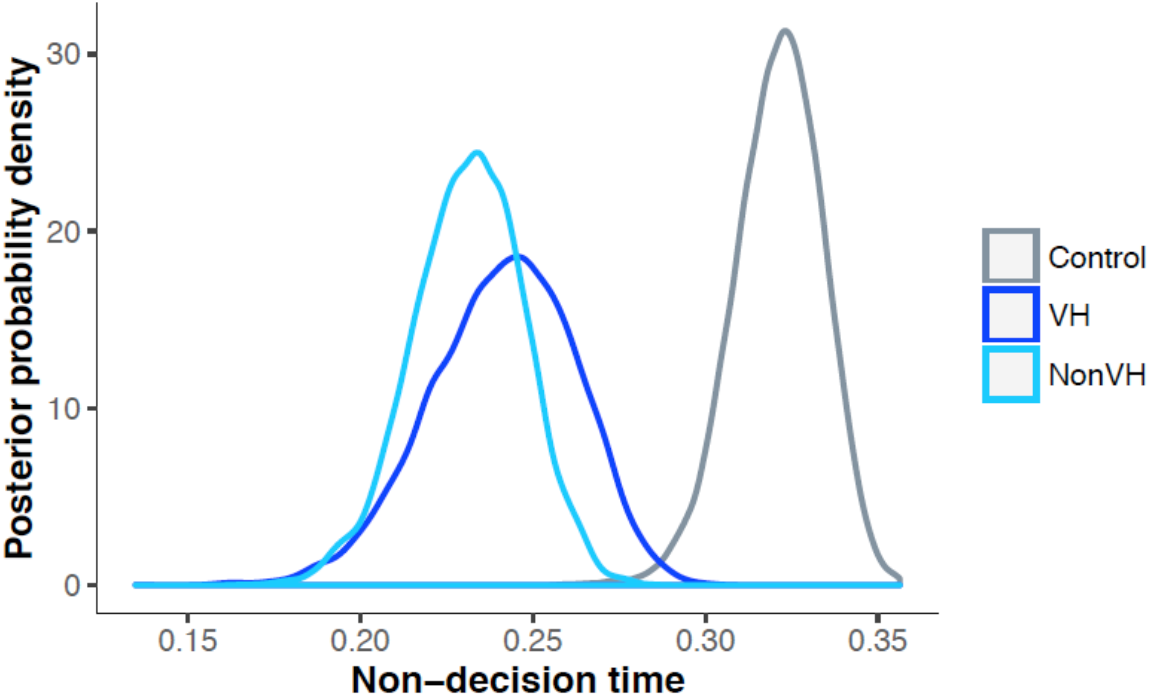
Non-decision time. ***Legend –*** Posterior probability density plots for non-decision time.

#### Comparisons between groups

*Comparisons across conditions*

Figure 5 shows posterior probabilities within each group for the three conditions (Controls: left panel; nonVH: middle panel; VH: right panel). For drift rates, controls showed an expected pattern with significantly longer drift rates in the incongruent condition compared to the neutral and congruent conditions (*P* = 0.28% and *P* = 0.32%), and similar rates in the neutral and congruent conditions (*P* = 50.92%). The nonVH group showed a similar pattern with significantly longer drift rates in the incongruent relative to the congruent condition (*P* = 0.15%), although not significantly different from neutral (*P* = 1.55%), with similar rates in the neutral and congruent conditions (*P* = 18.31%). In contrast, VH patients showed no significant differences between incongruent and neutral or congruent conditions (*P* = 3.28% and *P* = 17.47%), with similar rates also between neutral and congruent conditions (*P* = 17.80%).

**Figure 5.**
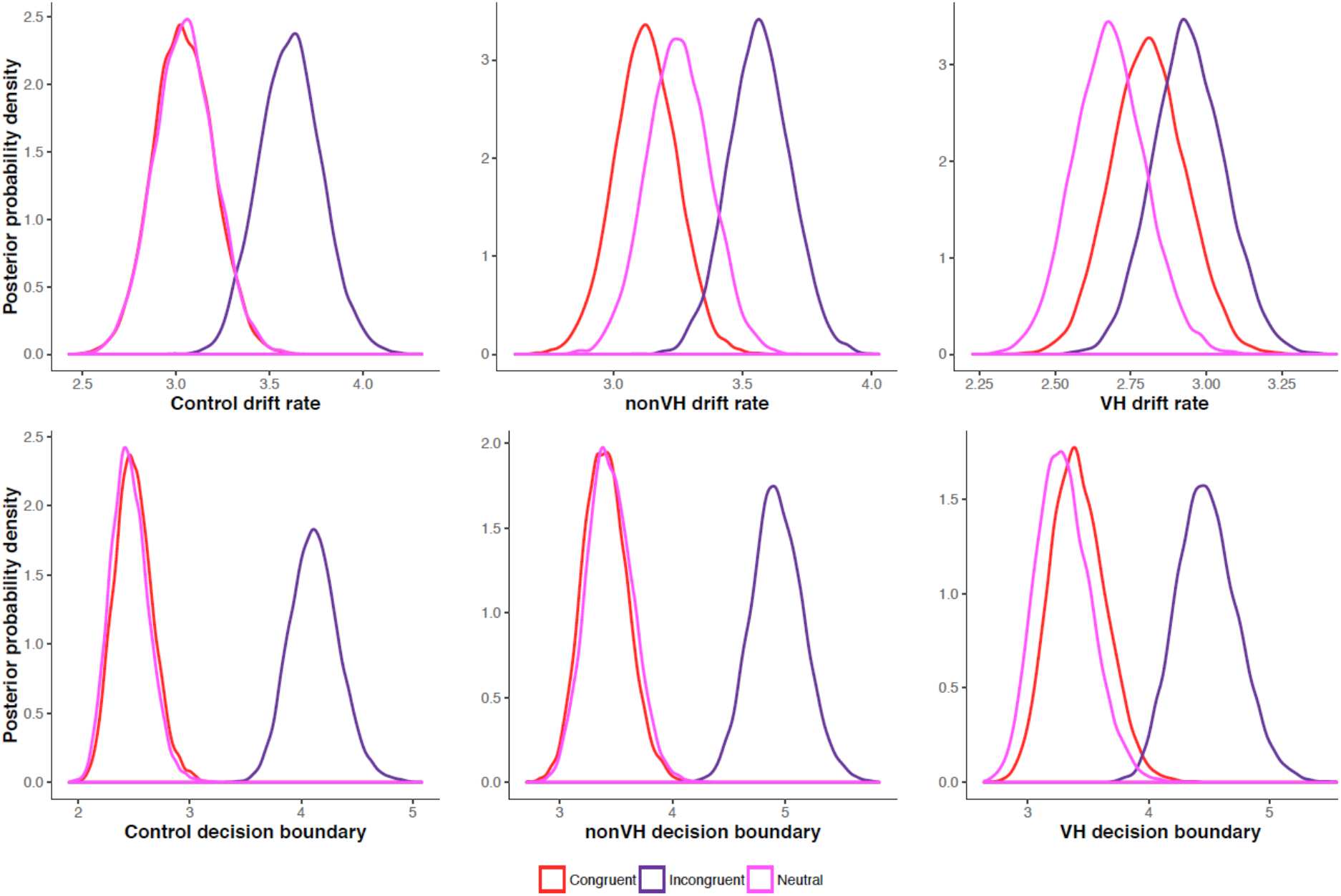
Within group comparisons of drift rates and decision boundaries. ***Legend –*** Posterior probability density plots for drift rates (top panel) and decision boundaries (bottom panel).

All groups showed a similar pattern for decision boundaries across the conditions, with significantly larger decision boundaries in the incongruent relative to the neutral and congruent conditions (Control: *P* = 0.00% and *P* = 0.00%; nonVH: *P* = 0.00% and *P* = 0.00%; VH: *P* = 0.00% and *P* = 0.00%), with similar thresholds in the neutral and congruent conditions (Control: *P* = 38.80%; nonVH: *P* = 37.32%; VH: *P* = 20.00%)

## Discussion

Our results demonstrate two features that characterise the perceptual decision making process of hallucinating patients relative to non-hallucinating patients: lower drift rates, and an inability to adjust drift rates to accommodate changes in perceptual conflict.

Behaviourally, all groups had 100% accuracy on the ANT. Employing a task with relatively low cognitive demands that PD patients could easily understand and execute was particularly important, given the possibility of cognitive impairment in this patient cohort. This ensured we could accurately access perceptual decision making without higher order cognitive deficits confounding performance. Hallucinating patients had the slowest reaction times on the ANT, although each group showed the expected pattern of shorter reaction times for the easier (neutral and congruent) conditions, and longer for the more difficult (incongruent) condition. Our results from the hDDM reveal the added benefits of modelling these data to uncover the cognitive processes underlying the behavioural results.

The drift rate parameter reflects how efficiently information is accumulated and determines the quality of evidence that enters the decision making process (Ratcliff and McKoon, 2008). In all three conditions, the control and non-hallucinating groups displayed similar drift rates. Hallucinating patients had consistently lower drift rates. In between-condition comparisons, drift rates are modulated by task difficulty (Voss *et al.*, 2004). This effect was apparent in our results for both controls and non-hallucinators. These groups had longer drift rates in the incongruent relative to the congruent conditions, suggesting their integration of information into the decision making process was flexibly modulated in response to task demands. In contrast, the hallucinating group had similar drift rates across the three conditions of perceptual conflict, indicating the absence of flexible context-dependent modulation of sensory accumulation.

Previous work has identified neural correlates of evidence accumulation during perceptual decision making. Recordings in monkeys suggest that neuronal populations in primary sensory areas (e.g., middle temporal visual area) fire in response to properties of a stimulus, and downstream regions (e.g., lateral intraparietal, frontal eye fields (FEFs), dorsal lateral prefrontal cortex (DLPFC)) integrate this information over time until sufficient evidence is accumulated for a decision (Newsome *et al.*, 1989, Gold and Shadlen, 2001, Shadlen and Newsome, 2001, Heekeren *et al.*, 2008). In humans, evidence accumulation has been related to frontoparietal regions (i.e., dorsal and ventrolateral PFC and FEF (Heekeren *et al.*, 2004, Heekeren *et al.*, 2006, Philiastides and Sajda, 2007, Liu and Pleskac, 2011)) and the anterior insula (Ho *et al.*, 2009), as well as integration across large-scale networks (Shine *et al.*, 2016). In PD, abnormal local and network-level engagement of frontoparietal and insula regions has been found in hallucinators (Stebbins *et al.*, 2004, Ramírez-Ruiz *et al.*, 2008, Goetz *et al.*, 2014, Shine *et al.*, 2015), and is typically equated with attentional dysfunction. In perceptual decision making, accumulation of sensory evidence and attention are highly collinear –and possibly inseparable– processes (Mulder *et al.*, 2014). It follows that our finding of impaired drift rates may offer a more tangible, computational framework to better specify attentional impairments in PD hallucinators.

All groups modulated their decision boundaries in response to the more difficult condition, displaying larger boundaries for the incongruent relative to congruent and neutral conditions. Flexible adaptation of decision thresholds in response to task demands has been shown previously in PD patients and related to a medial prefrontal cortex-subthalamic nucleus network that supports conflict detection and gating of decision thresholds when increased caution is required (Cavanagh *et al.*, 2011, Herz *et al.*, 2016b). Despite the flexible adaptation of thresholds, regardless of hallucination status, PD patients had larger decision boundaries than controls. The lack of difference between the hallucinating and non-hallucinating groups suggests that a widening of decision boundaries during perceptual decision making is a feature of PD more generally. This corresponds to findings in healthy ageing where older adults display wider decision boundaries relative to younger adults (Ratcliff *et al.*, 2001, Spaniol *et al.*, 2006, Ratcliff *et al.*, 2007). In older adults, adopting conservative decision criteria is presumed to be a compensatory strategy to prevent errors in speed/accuracy tradeoff tasks, and this effect may be amplified in PD.

Hallucinating and non-hallucinating patients also had comparable non-decision times, and these were shorter than controls. This might have seemed counter intuitive in light of the longer non-decision times that are found in ageing (Ratcliff, 2008, Starns and Ratcliff, 2010), but similar evidence of reduced non-decision times in PD has been shown with the application of an hDDM to a saccadic go/no-go task (Zhang *et al.*, 2016). Non-decision time encompasses diverse processes: encoding evidence from a stimulus and extracting its dimensions to guide the decision making process, and execution of a motor response (Ratcliff *et al.*, 2016). As response execution would not be anticipated to speed up in PD, decreased latencies in stimulus encoding and extraction of details may drive this finding. Future work may make this distinction in PD, although currently non-decision time is a relatively underspecified term compared to the other parameters in the DDM, and few studies have successfully separated its sub-components (Mulder *et al.*, 2014).

Inefficient and less flexible evidence accumulation during perceptual decision making emerges as a unique feature of PD hallucinators, whilst changes in cautiousness and visuo-motor processes are apparent regardless of hallucinator status. Longer drift rate latencies lead to slower decisions, and because perceptual decision making is a noisy process this increases the chance of errors (Krypotos *et al.*, 2015, Herz *et al.*, 2016a). In PD hallucinators, low drift rates that are invariant to changing environmental demands would be a source of low quality or inaccurate information entering the perceptual process.

Our results can be interpreted in light of models where an over weighting of top-down predictions, at the expense of bottom-up sensory information, contributes to hallucinations (Friston, 2005, Corlett *et al.*, 2009, Fletcher and Frith, 2009). Such models are formally described by a Bayesian framework: sensory input (bottom-up information, or likelihood) is integrated with known statistics about the environment (top-down predictions, or priors), forming an estimate of the external stimulus (the posterior). Contributions of the likelihood and prior in generating the posterior estimate are weighted in accordance with their certainty. Noisy sensory input is uncertain and carries less weight, shifting the balance in favour of priors. In this context, perception is vulnerable to excessive influence from internally generated beliefs and expectations. In PD, if visual information is accumulated slowly and inefficiently and is therefore less informative, this may contribute to the down weighting of bottom-up information in favour of top-down information.

Whilst the Bayesian framework is a compelling computational description of hallucinations, it does not necessarily favour a single mechanistic explanation (Teufel and Fletcher, 2016). For example, aberrant predictive coding across hierarchal brain circuitry (Friston *et al.*, 2014) and large scale disruptions in the brain’s excitatory-to-inhibitory (E/I) tone (Jardri *et al.*, 2017) both accommodate this account of hallucinations. Reconciling computational descriptions of hallucinations with a mechanistic framework will open important therapeutic avenues, as predictive coding and E/I accounts rely on distinct neuromodulatory and neurotransmitter profiles. In PD, we are beginning to uncover the psychological and neural signatures associated with a bottom-up vs. top-down imbalance in perception. Along with the results of the current study and previous evidence of lower level deficits in attention and visual processing, previous work has identified over activity in the default network of PD hallucinators (Franciotti *et al.*, 2015) and increased coupling between the default network and visual cortex (Shine *et al.*, 2015). Given the default network’s role in construction of mental imagery (Andrews-Hanna, 2012) and its positioning as a transmodal system distinct from unimodal sensory regions (Margulies *et al.*, 2016), over engagement of the default network during visual perception may be a source of excessive top-down influence. Future studies in PD will be needed to reconcile these insights with a mechanistic framework.

In summary, our results suggest that impaired drift rate can provide a novel computational framework encompassing the sensory processing and lower level attentional deficits previously described in visual hallucinators. Alterations in the dynamic process of evidence accumulation may therefore be a valuable marker to exploit in future explanatory and therapeutic studies of PD visual hallucinations.

## Acknowledgments

CO is supported by a National Health and Medical Research Council Neil Hamilton Fairley Fellowship (1091310). AJM is supported by an Australian Postgraduate Award through the University of Sydney. JMS is supported by a National Health and Medical Research Council CJ Martin Fellowship (1072403). SJGL is supported by a National Health and Medical Research Council Practitioner Fellowship (1003007)

